# Cryo-EM Structure of Bacterioferritin Nanocages Provides Insight into the Bio-mineralization of Ferritins

**DOI:** 10.1101/2021.02.04.429857

**Authors:** Chacko Jobichen, Tan Ying Chong, Rajesh Rattinam, Sandip Basak, Mahalashmi Srinivasan, Kannu Priya Pandey, Tran Bich Ngoc, Jian Shi, Jayaraman Angayarkanni, J Sivaraman

## Abstract

Iron is an essential element involved in various metabolic processes. The ferritin family of proteins forms nanocage assembly and are involved in iron oxidation, storage and mineralization. Although several structures of human ferritin and bacterioferritin subunits have been resolved, there is still no complete structure that shows both the trapped Fe-biomineral cluster along with the nanocage. Furthermore, whereas the mechanism of iron trafficking has been explained using various approaches, an atomic-level description of the pathway and the biomineralization that occurs inside the cavity are lacking. Here, we report three cryo-EM structures of different states of the *Streptomyces coelicolor* bacterioferritin nanocage (i.e., apo, holo) at 3.4 Å to 4.6 Å resolution and the subunit crystal structure at 2.6 Å resolution. The holo forms show different stages of Fe-biomineral accumulation inside the nanocage and suggest the possibility of a different Fe biomineral accumulation process. The cryo-EM map shows connections between the Fe-biomineral cluster and residues such as Thr157 and Lys42 from the protein shell, which are involved in iron transport. Mutation and truncation of the bacterioferritin residues involved in these connections can significantly reduce iron binding as compared with wild type bacterioferritin. Moreover, *S. coelicolor* bacterioferritin binds to various DNA fragments, similar to Dps (DNA-binding protein from starved cells) proteins. Collectively, our results represent a prototype for the ferritin nanocage, revealing insight into its biomineralization and the potential channel for ferritin-associated iron trafficking.

## Introduction

Iron is involved in various physiological processes, including DNA synthesis, gene regulation, respiration, and oxygen transport (Papanikolaou & Pantopoulos, 2005). However, free iron is toxic to cells, and is linked with the formation of free radicals from reactive oxygen species. Thus, iron levels are carefully regulated to prevent iron overload and oxidative stress. The ferritin family of proteins play an important role in stress management by sequestering excess iron (Fe^2+^) and storing it as insoluble Fe^3+^ clusters within a nanocage assembly. When required, these iron stores can be reduced and transported out of the cavity for cellular processes.

Ferritin-like molecules are present in archaea, bacteria, and eukarya, and their structures are well conserved despite poor sequence identity (Bradley, Le Brun et al., 2016). There are three types of ferritin-like molecules found in bacteria: bacterioferritin (Bfr), ferritin (Ftn), and DNA binding proteins from starved cells (Dps)(Smith, 2004). In contrast, humans have only one form, ferritin (Ftn). Bfr and Ftn nanocages are composed of 24 subunits and can store up to 4,300 Fe ions, whereas Dps nanocages comprise 12 subunits and able to store up to 500 Fe ions in the cavity(Bradley et al., 2016). Bfr proteins bind heme between two monomers at the dimeric interface, with the nanocage capable of binding 12 heme molecules. Fe^2+^ ions become trapped inside the ferroxidase center within the helical bundle of the Bfr monomer (Lawson, Treffry et al., 1989, Tosha, Ng et al., 2010).

Several crystal structures of human Ftn and Bfr have been resolved (Janowski, Auerbach-Nevo et al., 2008, Yao, Wang et al., 2012), and cryo-EM structures of mammalian Ftn nanocages have been reported, along with their functional studies (Ahn, Lee et al., 2018, Montemiglio, Testi et al., 2019, Russo & Passmore, 2014). Two groups have also reported biomineral core formation in ferritin (Pan, Sader et al., 2009, Pozzi, Ciambellotti et al., 2017). However, to date, there is no cryo-electron microscopy (cryo-EM) nanocage structure of Bfr. Moreover, none of the reported structures shows the three-dimensional structure of the nanocage cavity with the trapped Fe-biomineral cluster. Although the mechanism of iron trafficking has been detailed previously, an atomic-level description of the pathway and the process of biomineralization inside the cavity are still lacking.

*Streptomyces coelicolor* is known for its adaptability to environmental stress and its production of numerous antibiotics, immunosuppressants and anti-tumor agents. However, it does not cause disease in humans, plants, or animals (Bentley, Chater et al., 2002, Champness, Riggle et al., 1992, Hoskisson & van Wezel, 2019, Lakey, Lea et al., 1983, Lee, Karoonuthaisiri et al., 2005). Studies suggest that *S. coelicolor* employs a ferroxidase reaction to manage the stress generated by reactive oxygen species, which, in turn, results in the sequestration of iron within its nanocage (Lee et al., 2005). Thus, *S. coelicolor* is an ideal candidate for investigating the iron trafficking mechanism of ferritin.

Here, we report the high-resolution cryo-EM structures of the apo (empty) and two holo (containing iron-biomineral) states of the *S. coelicolor* Bfr (ScBfr) nanocage, along with the crystal structure of the ScBfr subunit. The holo-ScBfr structures (form-I and form-II) illustrate the various stages of mineralization of the Fe cluster sequestered inside the nanocage cavity. Notably, the holo-ScBfr form-II maintains connections between the protein shell and the Fe-biomineral cluster inside the cage, suggestive of potential pathways for iron accumulation and release. Additionally, we show that ScBfr binds to various DNA fragments and plasmids in vivo and in vitro. Taken together, this study helps to clarify the multiple functions of ferritin, particularly the molecular mechanism of iron transport.

## 2. Results

### 2.1 Cryo-EM structures of *Streptomyces coelicolor* bacterioferritin (ScBfr) nanocages

The 3D reconstructions of the ScBfr images show three major species of nanocages: the apo-ScBfr (no Fe-biomineral cluster) form, and two forms of holo-ScBfr—form-I and form-II— both with Fe-biomineral clusters accumulated/trapped in the centers of the cavities. The final reconstruction was built from ~77,000 particles for the apo-ScBfr nanocage, ~19,000 particles for the holo-ScBfr form-I nanocage, and ~13,000 for the holo-ScBfr form-II nanocage. We generated a 24-mer nanocage template model using our crystal structure of the ScBfr subunit (see section 2.3), and this model was fitted into the cryo-EM maps and used in the refinement. The apo-ScBfr and holo-ScBfr structures (I and II) had resolutions of 3.4 Å, 3.6 Å, and 4.6 Å, respectively, with imposed O symmetry (Fig 1A-C, See Supplementary Material fig S4A-C, Table 1).

**Figure 1:**
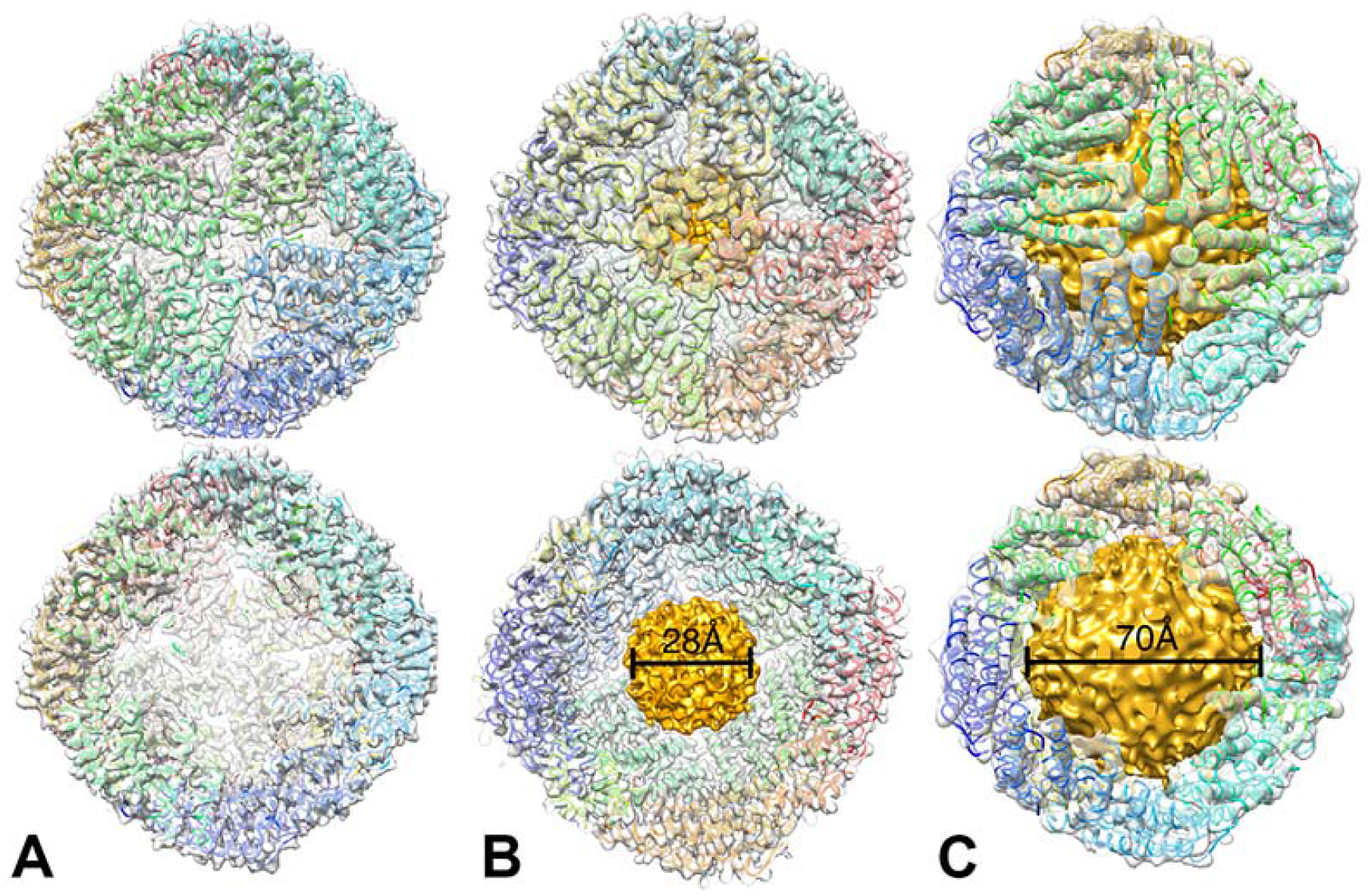
The cryo-EM structures of **(A)** apo-ScBfr and **(B)** holo-ScBfr-form-I and **(C)** holo-ScBfr-form-II. The trans-sectional view of the model shown below each form. The Fe-mineral accumulated in the nanocage cavity is shown in yellow color in holo-form-I and holo-form-II. The apo-ScBfr structure was solved at 3.4 Å resolution while the holo-form-I was solved at 3.6 and holo-form-II at 4.6 Å resolution. The holo-ScBfr form-II has a much larger Fe-biomineral cluster (diameter 70 Å) when compared to holo-ScBfr form-I (diameter 28 Å).

**Table 1:**
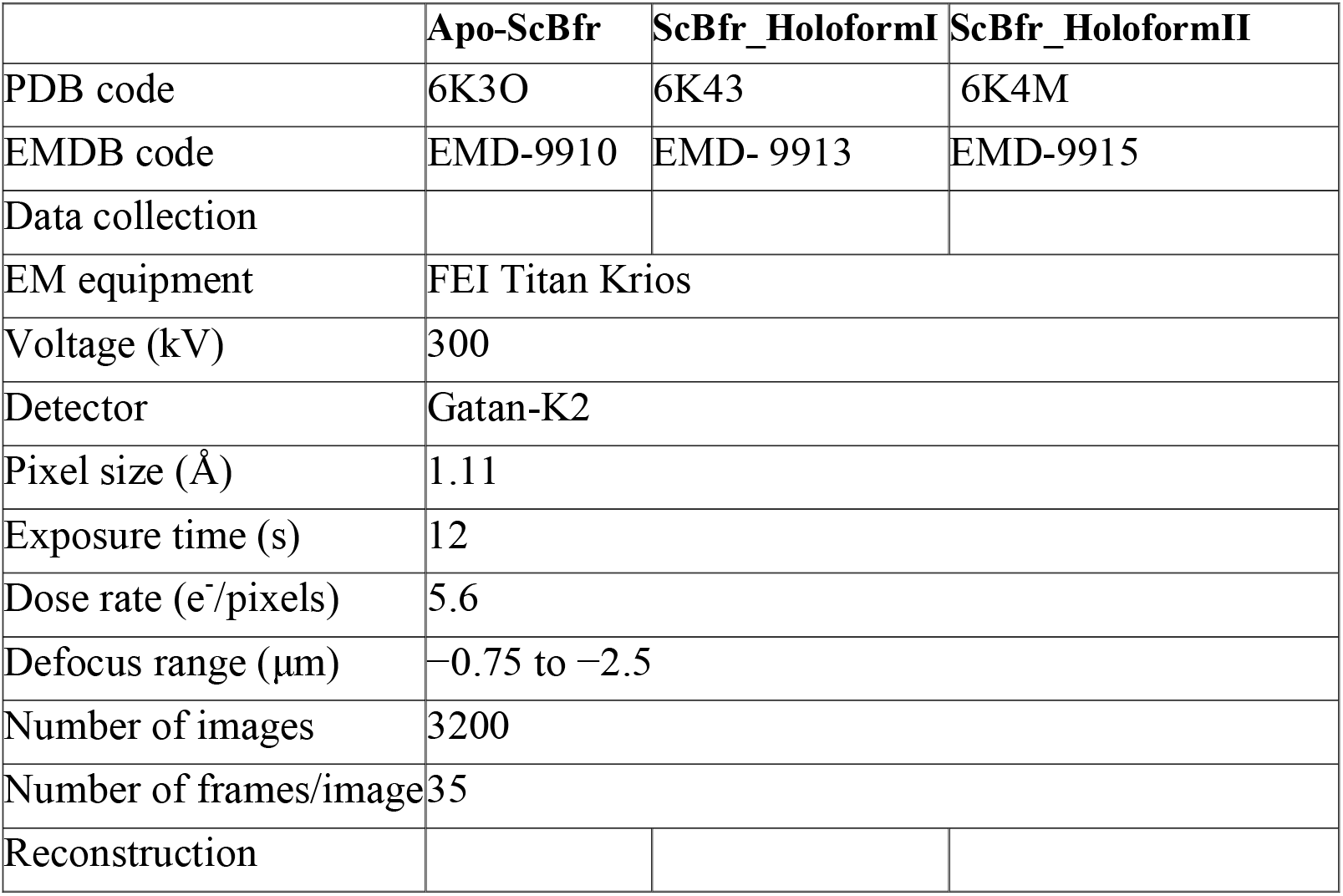
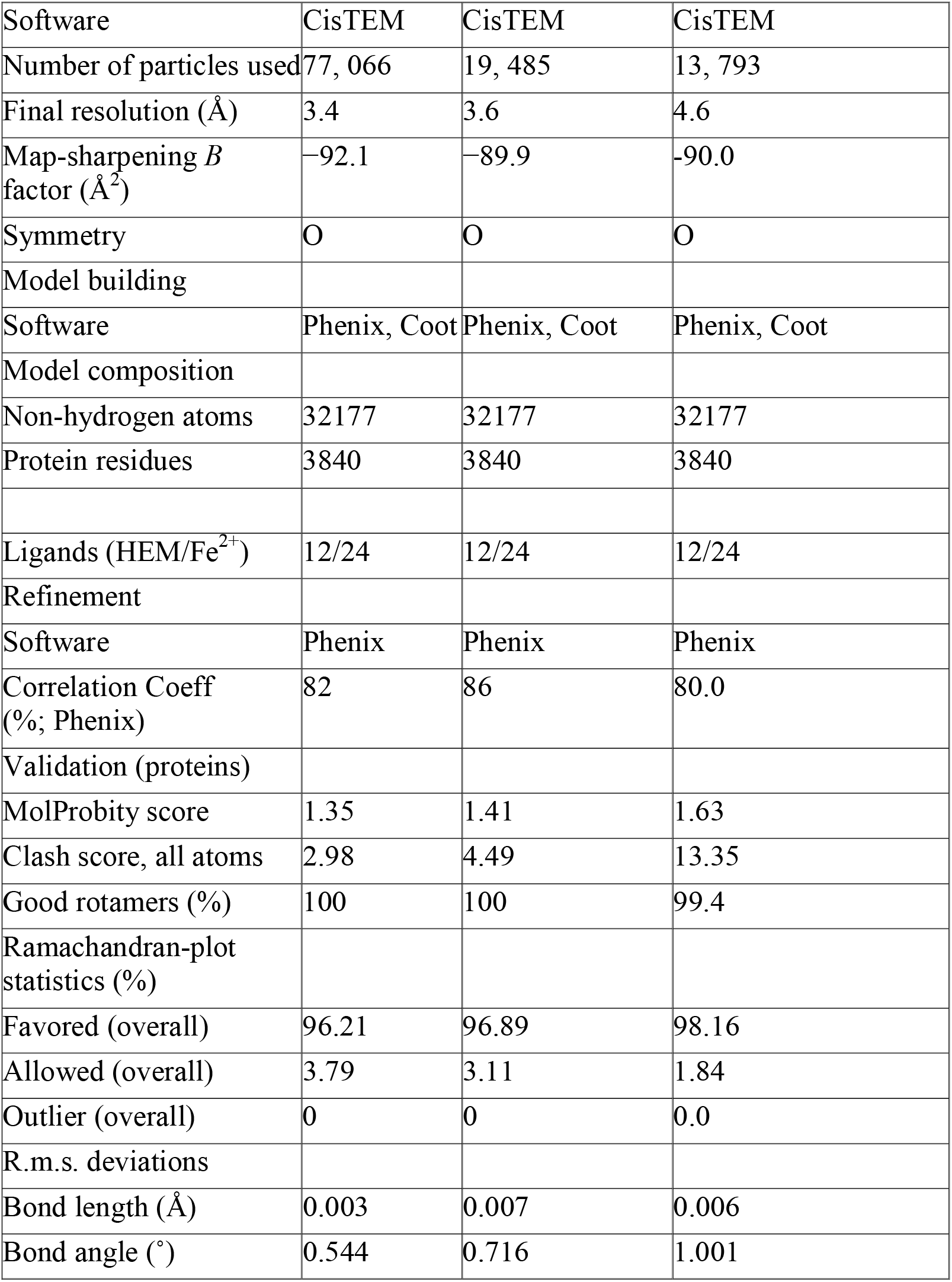
Cryo-EM data collection and processing

#### 2.1.1 Structure of apo-ScBfr nanocage

The apo-ScBfr nanocage has an external diameter of 125 Å and an internal diameter of 82 Å with an empty cavity (Fig 1A). The subunits in the nanocage are arranged in an anti-parallel fashion, forming a dimer (See Supplementary Material, fig S1A-B). These dimers are stacked perpendicular to the monomer from the second dimeric unit (See Supplementary Material, fig S1A). In total, 12 dimeric units assemble to form the nanocage. There are six symmetric 4-fold channels, eight symmetric 3-fold channels, and 24 B-pores formed by the subunits of the nanocage structure (See Supplementary Material, fig S1A). The 4-fold channels are formed by the C-terminal regions (146 to 157 aa) of the four monomers (one monomer from each dimeric unit), which protrudes toward the nanocage cavity (See Supplementary Material, fig S1A).

The cryo-EM map of apo-ScBfr (Fig 2A-C) shows density for all residues of the 24 chains of the ScBfr nanocage, except the last 10 residues in the C-terminus. The densities for the heme molecules are well defined in the dimeric interface of ScBfr (Fig 2B), and a total of 12 heme molecules are modelled in the nanocage. The Met52 residues from two monomers form coordination bonds with the heme molecule (Fig 2B), and residues Arg45, Phe49, and Glu56 from both monomers make hydrogen bonding contacts with heme. The density of the Fe^3+^ ion located in the ferroxidase center is also well defined, and residues Glu18, Glu51, His54, Glu94, Glu127, and His130 form coordination bonds with the Fe^3+^ ion (Fig 2C). Further, self-assembly of the subunits in the nanocage is predominantly maintained by electrostatic interactions (See Supplementary Material, fig S1C).

**Figure 2.**
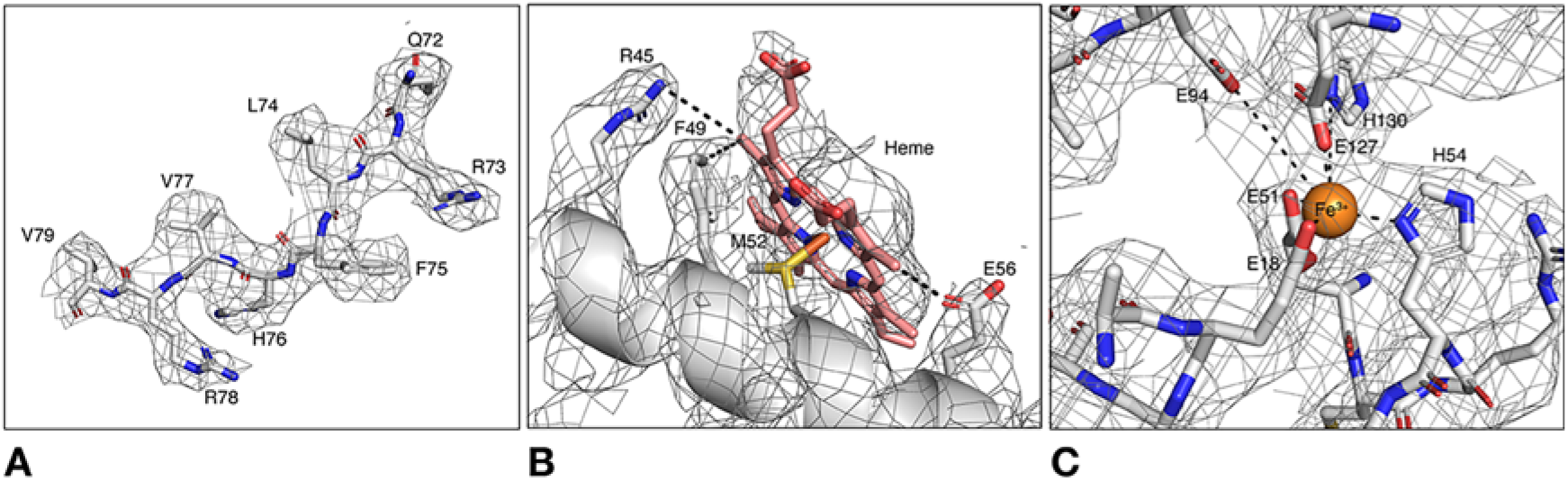
**A:** The representative cryo-EM map of apo-ScBfr nanocage to show the quality of the map. **B**: The cryo-EM map of apo-ScBfr nanocage showing the heme molecule and the adjacent residues. The heme molecule is sandwiched between the residue Met52 from both monomers of the dimer, for clarity, only Met52 from one monomer is shown in this figure. **C**: The density map of apo-ScBfr nanocage showing Fe ion in the ferroxidase center and the coordinating residues.

#### 2.1.2 Structure of holo-ScBfr nanocages (form-I and form-II)

The Fe-biomineral–trapped holo-ScBfr nanocage structures have an internal diameter of 82 Å and an external diameter of 125 Å, similar to the apo-ScBfr nanocage. There were two models with Fe-biomineral accumulation/deposition inside the nanocage cavity: form-I has a smaller diameter of Fe-biomineral (28 Å) inside the nanocage (Fig 1B), whereas form-II has a much larger diameter of Fe-biomineral (70 Å) (Fig 1C) at 1.6-σ contour level.

The Fe-biomineral volume is 5% of the total volume of the nanocage cavity in form-I but 70% in form-II. The Fe-biomineral core inside the nanocages of both forms I and II do not fit to any symmetry. Consistently, others (Bradley et al., 2016, Crabb, Moore et al., 2010, Theil, Tosha et al., 2016) have suggested that the chemical content and structure of iron inside the core is polyphasic or heterogenous and may not be packed regularly. Moreover, the enhanced deposition of Fe-biomineral inside the cavity is presumably due to protein overexpression in the recombinant bacteria, and that this stressful environment facilitates the active conversion of Fe^2+^ to Fe^3+^ (Guerrero Montero, Dolata et al., 2019).

### 2.2 Insights into the pathway of Fe ion movement

Although various pathways (Bradley et al., 2016, Bradley, Moore et al., 2017a, Bradley, Svistunenko et al., 2017b, Yao et al., 2012) have been proposed to explain the passage of iron in and out of the nanocage cavity, the exact mechanism remains unclear. In this context, our structures of the holo forms of the ScBfr nanocages reveal three intriguing features.

First, in the holo-ScBfr form-II, the cryo-EM map of the nanocage structure shows connections between the trapped Fe-biomineral inside the nanocage cavity and residue Thr157 in the C-terminus of each monomer, forming a continuous pathway spanning 38 Å between the ferroxidase center in each monomer and the nanocage cavity (Fig 3A, 3B). This connection represents one possible route for Fe^3+^ ion transport from the ferroxidase center into the nanocage cavity. Turano *et al* (Turano, Lalli et al., 2010) proposed a similar pathway based on NMR studies of the *Lithobates catesbeiana* M-ferritin subunit. Superposition of the monomeric ScBfr (crystal structure, see section 2.3) with the *L. catesbeiana* M-ferritin subunit showed similarity in at least 12 residues (Ala21, Tyr25, His28, Glu47, Val83, Phe87, Asp90, Leu134, Leu138, Glu146, Glu155, and Thr157) involved in the movement of Fe^3+^ from the ferroxidase center to the nanocage cavity (Fig 3B). Notably, most of these residues are highly conserved among Bfr proteins (See Supplementary Material, fig S2A) and we also observed a continuous negative patch between the ferroxidase center and the C-terminal (See Supplementary Material, fig S1C). To confirm the role of these residues, in particular the C-terminal region, in iron transport, we generated a C-terminal truncated version of Bfr (ScBfr_1-152_: residues 153-167 removed) and examined Fe binding compared with the wild type. We observed significantly reduced Fe binding in the C-terminal truncated mutant as compared with the wild type (0.04 ppm vs. 0.27 ppm), suggesting an important role for this C-terminal region in Fe-biomineral accumulation.

**Figure 3:**
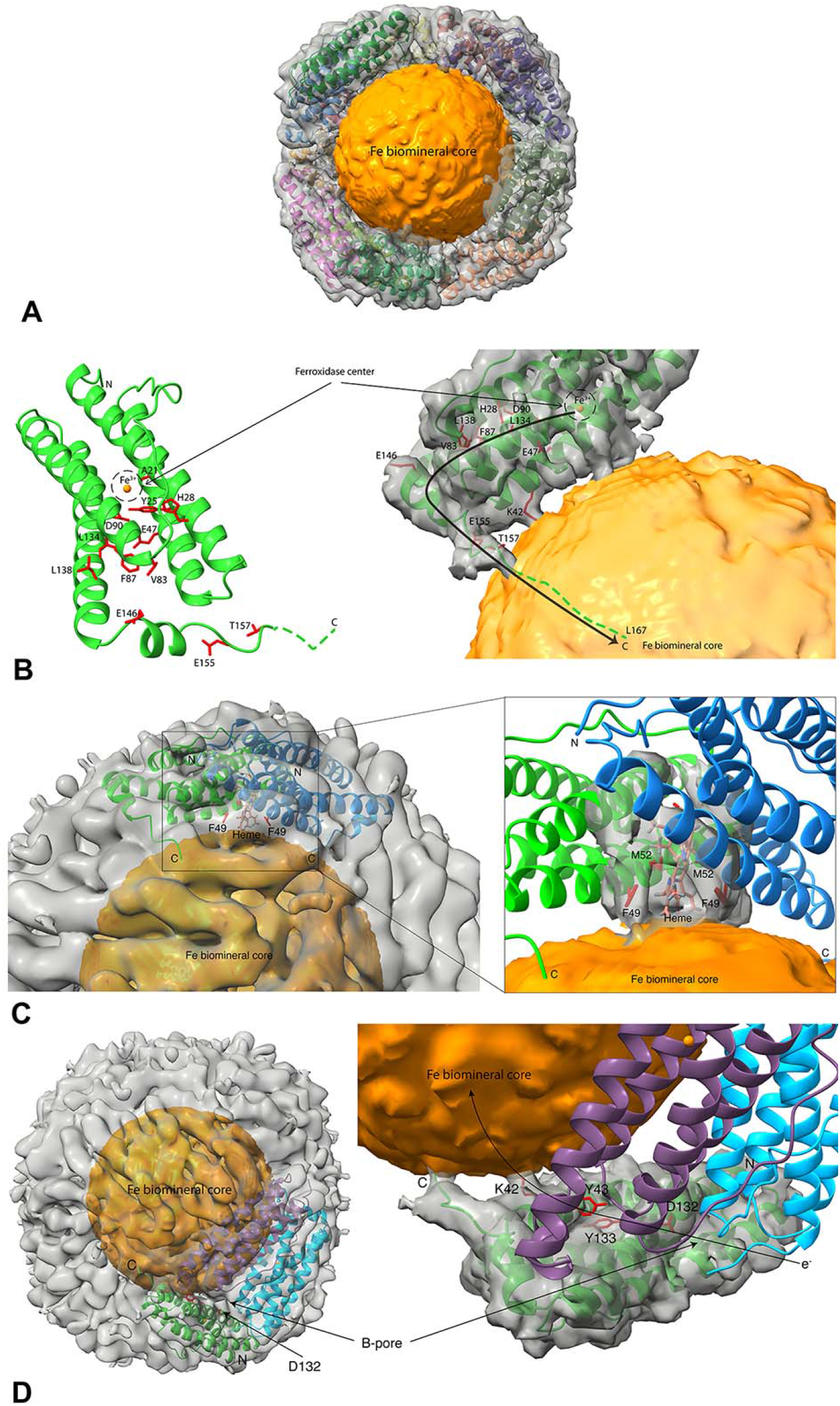
**A:** Overall view of the ScBfr nanocage with the trapped Fe-biomineral cluster. **B:** The density map of the nanocage showing the connections between the trapped Fe-biomineral cluster inside the nanocage cavity and the ScBfr protein shell. **(B-right panel):** The Thr157 in the C-terminal of each monomer is connected to the Fe-core in the nanocage cavity and thus forms a continuous pathway spanning 38Å between the ferroxidase center located in each monomer and the nanocage cavity (shown as black arrow). **(B-left panel):** The 12 residues possibly involved in the movement of Fe^3+^ from the ferroxidase center to the nanocage cavity in ScBfr is shown as stick model and labelled. Fe atoms are shown as orange spheres. **(C):** The heme molecule sandwiched between the dimeric interface is also having connections with the Fe-core through the adjacent residues Met52 and Phe49. **(D):** Residues such as Lys42 and Tyr43 structurally located near the C-terminal region of ScBfr, along with Tyr133 and Asp132 connect the B-pore at the surface to the Fe cluster in the nanocage cavity. This figure was prepared using Chimera program(Pettersen, Goddard et al., 2004).

Second, in the cryo-EM map, we noted another connection between the Fe-biomineral and the heme binding region of the holo-ScBfr form-II, involving residues Met52 and Phe49 (Fig 3C). Whereas Met52 is highly conserved in Bfr proteins, Phe49 is conserved only in *S. coelicolor* and *Mycobacterium tuberculosis*. Previous studies in *E.coli* showed that heme plays an important role in the release of iron from the central cavity of the Bfr nanocage by facilitating electron transfer, and that reduction of Fe^3+^ to Fe^2+^ is the first step in the release of Fe stored in the Ftn nanocage (Yasmin, Andrews et al., 2011). In the present study, the observed connection in holo-ScBfr form-II suggests the possibility that the electrons transferred by heme enter the nanocage cavity to facilitate the conversion of Fe^3+^ to Fe^2+^ and aid in the subsequent release of Fe^2+^ from the cavity (Fig 3C). It is worth mentioning that a similar heme-mediated electron transfer process has been described in the Bfr-B–Ferredoxin (BfrB-Bfd) complex structure (PDB: 4E6K) (Yao et al., 2012).

Third, we also observed a continuous connection between the B-pore at the surface and the Fe-biomineral in the nanocage cavity. This is achieved between Lys42 and Tyr43, which are structurally located near the C-terminal region of ScBfr, and residues Tyr133 and Asp132. Residue Lys42 is connected to the Fe-biomineral core, whereas Asp132 is connected to the B-pore located on the surface of the nanocage (Fig 3D).

We speculate that this connection might play a role in electron transfer into the nanocage cavity as well as Fe-ion transfer. Indeed, mutating this conserved Lys42 to Ala (ScBfr K42A) reduces Fe binding as compared with the wild type (0.06 ppm vs. 0.27 ppm). Notably, in *E. coli* Bfr, Tyr34 has been reported to act as a molecular capacitor/electron donor for facilitating Fe oxidation (Bradley et al., 2017b, Ebrahimi, Hagedoorn et al., 2013) and Trp133 facilitates electron transfer (Bradley et al., 2017b). Collectively, these findings suggest that Tyr43 and Tyr133 in ScBfr might also contribute to electron transfer.

### 2.3 Crystal structure of ScBfr and nanocage model

The crystal structure of ScBfr was determined at 2.6 Å resolution (See Supplementary Material, Table 2). The structure is well defined in the electron density map, except for the last five residues (Gly163-Leu167), which are disordered and hence not modelled. The monomeric ScBfr adopts an α-helical structure with 4 α-helices bundled together followed by a short 2-turn helix at the C-terminus (See Supplementary Material, fig S2B), similar to its homologs (Khare, Gupta et al., 2011). As with our observations of the apo-ScBfr nanocage cryo-EM structure, each monomer is bound with one Fe ion in the ferroxidase center located within the helical bundle (See Supplementary Material, fig S2B). Four glutamic acids (Glu18, Glu51, Glu127, Glu94-water mediated) and His54 from 3 helices are involved in the Fe coordination in the ferroxidase centre (See Supplementary Material, fig S2B). Previous reports of Bfr crystal structures suggest that Bfr can form various crystal packing depending on the space group (i.e., 1 molecule to 48 molecules in the asymmetric unit) (See Supplementary Material, Table S2); yet, no Fe-biomineral has been observed in any of these asymmetric unit oligomers.

**Table 2:** Crystallographic data collection and refinement.

|  | **ScBfr** |
| --- | --- |
| **Data Collection** |  |
| Space group | I4 |
| Cell Dimensions |  |
| a,b,c (Å) | 123.76 123.76 172.30 |
| $\alpha, \beta, \gamma (^{\circ})$ | 90 90 90 |
| Resolution (Å) | 43.8-2.6 (2.69-2.60) |
| R<sub>merge</sub> | 0.05(0.15) |
| I/ σI | 3.1 (1.3) |
| Completeness (%)<sup>*</sup> | 97 (97) |
| Redundancy<sup>*</sup> | 5.0 (3.2) |
| **Refinement** |  |
| Resolution (Å) | 43.8-2.6 (2.63-2.6) |
| No. reflections | 76305 (2059) |
| R<sub>work</sub>/ R<sub>free</sub> | 0.161/0.208 |
| No. atoms |  |
| Proteins | 7817 |
| Ligand/Ion | 12 |
| Water | 625 |
| B-factors(Å<sup>2</sup>) |  |
| Proteins | 18.8 |
| Ligand/Ion | 55.58 |
| Water | 23.3 |
| R.m.s deviations |  |
| Bond lengths (Å) | 0.007 |
| Bond angles (°) | 0.899 |
| PDB Code | 5XX9 |

<sup>\*</sup>Values for the highest-resolution shell are given in parentheses.

A structural homology search for ScBfr identified conservation among several ferritins and Dps proteins (See Supplementary Material, Table S3). ScBfr has homologs in over 100 species, including pathogenic *M. tuberculosis* and *Corynebacterium diphtheria.* A comparison of the *M. tuberculosis* Bfr structure showed an overall similarity with the ScBfr structure (See Supplementary Material, fig S2C). The C-terminal regions of human H-Ftn, human L-Ftn, and bacterial Ftn (PDB code; 2FHA, 2FFX, 4XGS) are well ordered with an α-helical secondary structure that folds back to the core protein (See Supplementary Material, fig S2D). In contrast, the C-terminal tail of ScBfr (up to residue 162 present in the crystal structure) is flexible and can be extended up to 23 Å towards the center of the nanocage; unlike in human and bacterial Ftn, the ScBfr C-terminal tail might reach the center of the cavity (Fig 3B). The flexible C-terminal region of *S. coelicolor* might increase iron chelation into the core of the nanocage; indeed, *S. coelicolor* is known for its adaptability to environmental stress (Lee et al., 2005). A flexible/extended C-terminus has previously been implicated in ferroxidase activity and iron release in *M. tuberculosis* BfrB (Khare et al., 2011). Indeed, C-terminal truncation of Bfr significantly reduced Fe binding as compared with wild-type Bfr (see section 2.2).

In our homology search, Bfr and Dps proteins are well aligned in the 4-helical bundle core region (DpsA seq. identity: 22%; rmsd: 1.3 Å for 124 Cα atoms and DpsC seq. identity: 12%; rmsd: of 2.5 Å for 127 Cα atoms) (See Supplementary Material, fig S3A-B). However, residues 136-162 aa of the C-terminal region of Bfr (part of which connects the ferroxidase center and the center of the nanocage; please refer section 2.2) do not align with DpsA. Notably, Dps nanocages are formed by 12-mers; this differs from Bfr, which forms 24-mer nanocages. Besides, the ferroxidase center of Bfr is inside the 4-helical bundle of the monomer but located in the dimeric interface of Dps proteins (Ren, Tibbelin et al., 2003).

Using our crystal structure, we generated ScBfr nanocage model. A comparison of this model with the cryo-EM apo / holo ScBfr nanocages indicates that Cα atoms must move ~2.5 Å to 3.5 Å in various places to fit with the cryo-EM nanocage structures. Moreover, comparing the human Ftn nanocage cryo-EM structures with that of ScBfr shows a difference of ~3 Å to 5 Å in various regions (See Supplementary Material fig S2E). Notably, the C-terminal tail at the entrance of the nanocage cavity shows more flexibility for ScBfr than human Ftn, presumably because it is flexible and protruding out in ScBfr.

### 2.4 ScBfr binds with DNA

Dps proteins form stable contacts with DNA, irrespective of sequence, size or topology (Hitchings, Townsend et al., 2014). Notably, Dps proteins bind with chromosomes non-specifically and form complexes in which chromosomal DNA is condensed and protected from a diverse range of damage (Ceci, Cellai et al., 2004, Ceci, Mangiarotti et al., 2007). Recent studies have indicated that Bfr from *M. tuberculosis* and *Desulfovibrio vulgaris* can bind to plasmid DNA (Mohanty, Subhadarshanee et al., 2019, Timoteo, Guilherme et al., 2012). Given the structural similarities between ScBfr and the Dps proteins, we carried out electrophoretic mobility shift assay (EMSA) studies to ascertain whether ScBfr similarly binds DNA. We found PET32 plasmid binding to ScBfr, and also showed that DNA fragments co-purified with the ScBfr protein (See Supplementary Material, fig S6). Through DNA sequencing, we found that these co-purified DNA fragments are from the expression vector as well as from the genomic DNA of the host (See Supplementary Material, Table S4). Thus, ScBfr may have a possible role in DNA protection.

## 3. Discussion

The iron-storage function of ferritin plays a central role in iron homeostasis in the cell (Anderson & Frazer, 2017). Despite the wealth of structural and functional data available for ferritin-like molecules, the mechanism of Fe-biomineral accumulation inside the nanocage and its subsequent release for cellular activities remains unclear. In Bfr, Fe ions can be found inside the ferroxidase center, bound to heme in between monomers (i.e., at the dimeric interface), and as an Fe-biomineral cluster inside the nanocage cavity. In our cryo-EM structures of the ScBfr nanocage, we show Fe ions in all three of these locations. Notably, we show a physiologically relevant structure of a 24-mer ferritin nanocage with a trapped Fe-biomineral. In the holoform-II cryo-EM structure, the C-terminus of ScBfr is linked to the Fe-biomineral within the nanocage cavity (Fig 3B). In addition, the heme that is sandwiched between two monomers of ScBfr has a connection with the Fe-biomineral in the cavity through Phe49 (Fig 3C).

Using scanning transmission electron microscopy images from human liver samples, Pan *et al* (Pan et al., 2009) proposed that the ferric oxide biomineral can grow from the inner surface of the nanocage towards the center of the core. Similarly, Pozzi *et al* (Pozzi et al., 2017) reported that the biomineral core grows from the inner surface of the protein nanocage. This study was performed by soaking human L-Ftn crystals with ferrous solutions and determining the structure of the L-Ftn monomeric subunit at various time points (PDB 5LG8, 5LG2). The holo form-II model from our study shows that the EM density corresponding to the Fe-biomineral core does not interact with the inner surface of the protein, even at low contour levels (~1.1 σ) except the specific connections mentioned previously (refer section 2.2, Fig 3B-D, 4A). At high contour levels (~3.6σ), the density is present only at the center of the core (Fig 4A); this is supported by the unbiased 2D classes (Fig 4B). This observation tempting to suggest that the Fe biomineral accumulation might originate from the center of the nanocage as one among the possibilities.

**Figure 4.**
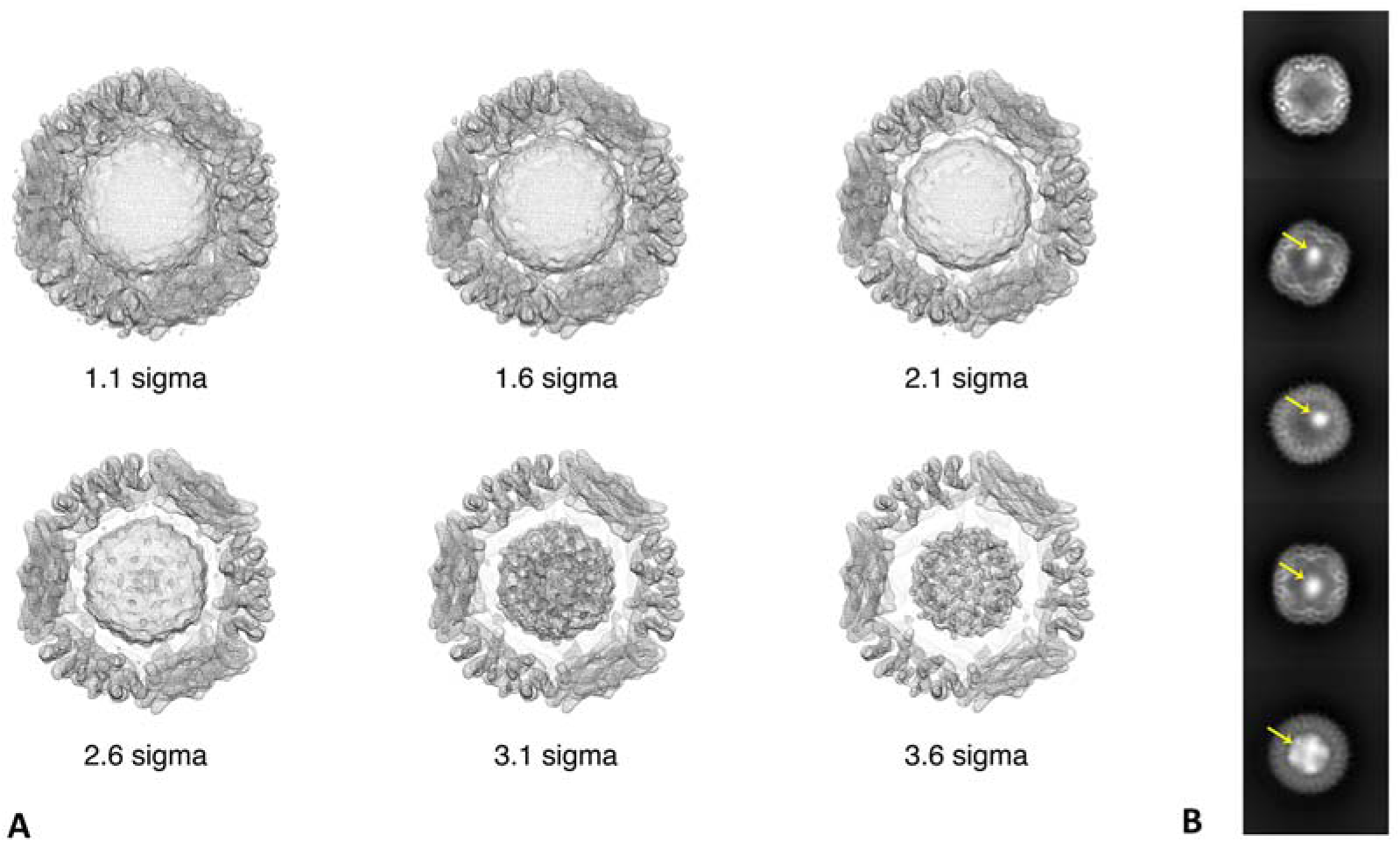
**(A)**. The cryo-EM density map of the holo form-II model at various contour level (from 1.1 to 3.6 sigma). The EM density corresponding to the Fe-biomineral core is not interacting with the inner surface of the protein even at low contour level (~1.1 sigma) except the specific connections mentioned in section 2.2; while at high contour level (~3.6 sigma) the density is present only at the center of the core and this is supported by the 2D classes (Fig. 4B). However, we observed clear density for the various connections between the Fe-biomineral core and the inner surface of the nanocage even at a contour level of ~1.6 sigma. Also refer to the figure 3B-D for various connections between the Fe-biomineral core and the protein shell. This figure was prepared using Chimera program (Pettersen et al., 2004). **(B)** The unbiased 2D classes obtained from the ScBfr particles. The yellow arrow points to the biomineral core observed in these classes.

Further we observed that the flexible C-terminal region of ScBfr may extend towards the center of the nanocage cavity (Fig 3B). The role of the C-terminal in the Fe-oxidation and reduction process is well documented in bacterioferritin (Khare et al., 2011). We also show that, C-terminal truncation of ScBfr significantly reduced iron binding as compared with wild-type. Comparatively, in human L-Ftn structures, the C-terminal region maintains a proper helical structure and is folded back to the inner surface of the nanocage shell (See Supplementary Material, fig S2C-E). It is worth mentioning that Bfr proteins have heme-binding properties and they also possess a ferroxidase center (Ebrahimi, Hagedoorn et al., 2015); neither of these features are found in L-Ftn (Ebrahimi et al., 2015). Moreover, in our study, the Fe-biomineral was co-purified with Bfr, whereas, in the L-Ftn study (Pozzi et al., 2017), it was incorporated through soaking. Collectively, these observations point to the likelihood of different mechanisms of Fe-biomineral accumulation in ferritins.

Combining our observations with those of previous literature (Bradley et al., 2017b, Ebrahimi et al., 2013, Rui, Rivera et al., 2012, Turano et al., 2010, Yasmin et al., 2011), we propose the following mechanism to explain Fe trafficking in and out of the nanocage cavity (Fig 5). Fe^2+^ ions enter through the B-pore and move into the ferroxidase center, where Fe^2+^ is converted to Fe^3+^ (Rui et al., 2012). These Fe^3+^ ions then move through the 4-helical bundle channel to reach the C-terminal helix of ScBfr (Fig 3B), and undergoes the processes of olation/oxolation that give rise to the ferrihydrite mineral core in the nanocage cavity (Bradley et al., 2016, Pettersen et al., 2004, Rivera, 2017). For outward trafficking, we propose that heme is involved in transferring electrons into the nanocage cavity (Fig 3C). Using these electrons, Fe^3+^ is reduced to Fe^2+^, with Fe^2+^ then exiting either through the B-pore or via attachment to a chelating agent (Yao et al., 2012, Yasmin et al., 2011).

**Figure 5:**
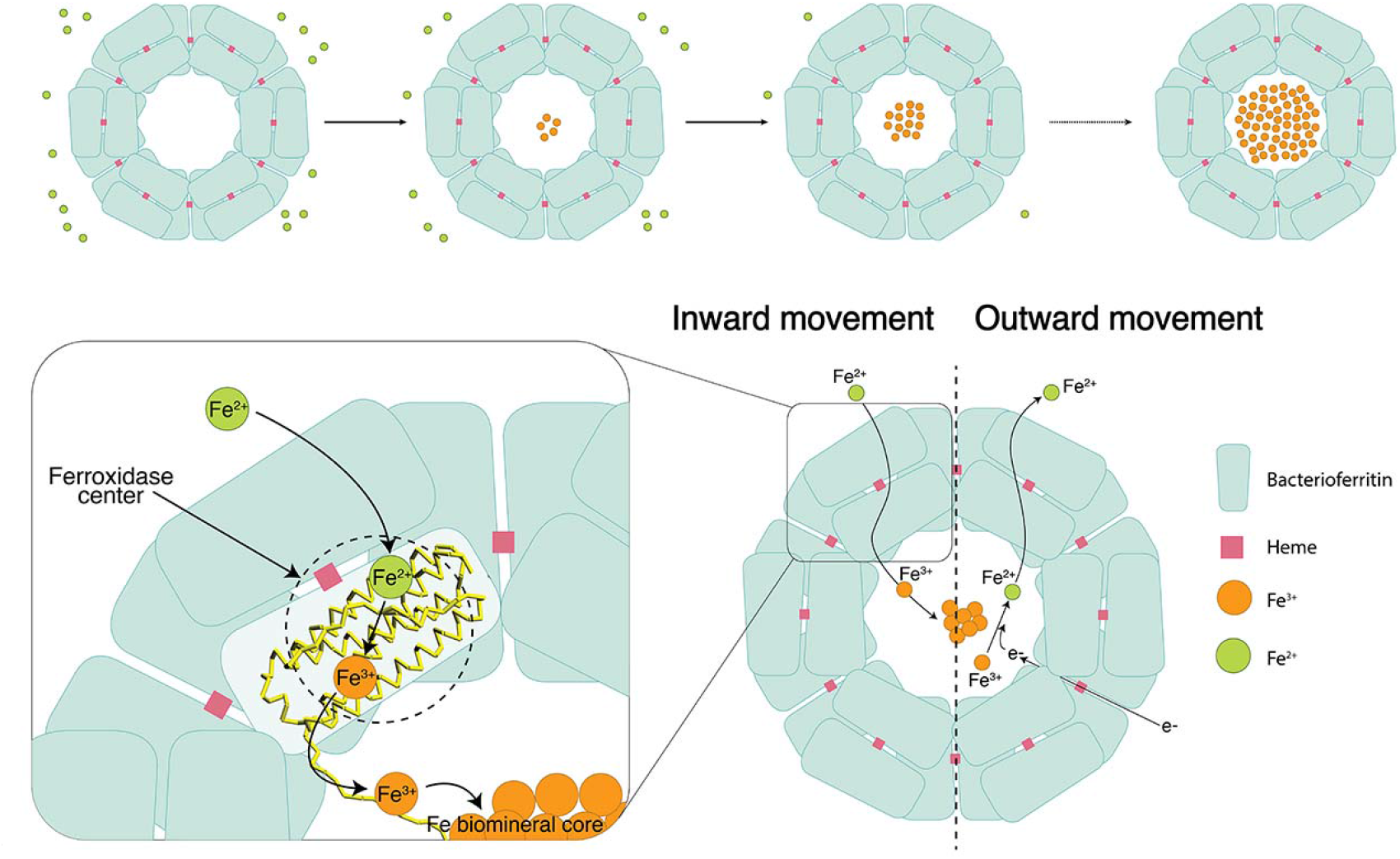
Schematic representation showing the inward and outward movement of iron in bacterioferritin nanocage. The Fe^2+^ atom enters into the Bfr shell through the B-pore and move to the ferroxidase center where it gets oxidized to form Fe^3+^. Fe^3+^ moves through the four-helix bundle into the C-terminal region of the Bfr and gets detached from the C-terminal and enters into nanocage cavity for storage. The flexible C-terminal region of ScBfr can extend up to the center of the nanocage cavity, and this might be crucial for nucleation initiation. Similarly, heme molecule will help to transport electron (e−) into the nanocage cavity, once the Fe^3+^ is reduced it is converted to Fe^2+^ that can move out of the Bfr shell.

A notable feature of this study is the three forms of ferritin nanocage structure: apo, holo I and holo II. Since the Fe-biomineral accumulation is a dynamic process (Ueno, Abe et al., 2009), we believe that there are various intermediate forms of the nanocage structures other than these three reported structures, and it is supported by the observation of several intermediates in the 2D classes (Fig 4B). We hypothesize that only these three forms (apo, holoform I and holoform II) have a sufficient number of particles to model a complete structure at high resolution. This could be because these intermediates (holo form I and II) are more stable than other intermediates, allowing more particles to be accumulated (Fig 1B-C).

Dps proteins oxidize and deposit iron in the form of ferric oxide and can bind DNA (Hitchings et al., 2014). Here, we show that ScBfr can also bind with DNA, suggesting that Bfr proteins can perform similar functions to Dps proteins (Mohanty et al., 2019, Timoteo et al., 2012). Although the exact role of Bfr proteins in DNA binding needs to be explored, we hypothesize that Bfr proteins might be able to replace or carry out the function of Dps proteins in species without Dps proteins; for example, in the case of *M. tuberculosis* (Mohanty et al., 2019).

In summary, here we report the apo and holo states of ScBfr nanocages with Fe ions, heme molecules, and Fe-biomineral clusters. The holo forms show different stages of Fe-biomineral accumulation inside the nanocage cavity. In particular, the holo form-II cryo-EM map shows various connections between the Fe-biomineral cluster and several residues of the protein shell. These residues are involved in iron transport and are hence critical for the movement of Fe ions in and out of the nanocage cavity. This study provides insight into the possibility of different mineralization process in bacterioferritin. These ScBfr nanocage structures are the representative structures of bacterioferritin family of proteins with Fe-biomineral trapped nanocage. Moreover, we show that ScBfr binds to DNA fragments in vivo and in vitro, suggestive of a possible role for ScBfr in DNA protection, a function previously ascribed to Dps proteins. Taken together, this study helps to clarify the multiple functions of ferritin, particularly the molecular mechanism of iron mineralization and transport.

## Materials and methods

### Cloning expression and protein purification

Bacterioferritin from *Streptomyces coelicolor* (ScBfr) was cloned into expression vector pET22b (Gene-Script, USA) and the recombinant plasmid was transformed into *Escherichia coli* BL21 competent cells for protein over expression. The transformed BL21 cells were used to inoculate LB medium (1L) containing (100 mg/ml) Ampicillin and the culture was grown at 37°C. After reaching the OD at A_600nm_ of 0.6-0.8, the culture was induced by the addition of 0.3 mM IPTG followed by incubation at 28°C for 7 hr. Cells were harvested by centrifugation at 6000 rpm using a JLA8.1 rotor in a Beckman Coulter centrifuge for 30 min at 4°C. The harvested cells were resuspended in 30 ml of lysis buffer (30 mM Tris pH 7.0, 200 mM NaCl, 1 mM PMSF, 5% Glycerol, 0.5% Triton x100) and lysed in sonicator. The resulting lysate was centrifuged at 18000 rpm (JA20 rotor, Beckman coulter) for 30 min at 4°C. The supernatant was added to Ni-Agarose affinity column equilibrated with same buffer. The unbound proteins were washed off with the same buffer thrice. The protein was then eluted twice with increased imidazole concentration of 250 mM and 350 mM. The eluted protein samples were injected into Superdex S-75 column equilibrated with gel filtration buffer (30 mM Tris pH 7.0, 200 mM NaCl, and 5% glycerol). The peak fractions were pooled and concentrated and used for cryo-EM and crystallography experiments. The purity and homogeneity of ScBfr was verified by using dynamic light scattering (DLS) experiments. DLS experiment indicated a polydispersity index of 0.30. Moreover, using the above described protocol, we have expressed and purified two mutant versions such as ScBfr K42A and ScBfr_1-152_ for the Inductively Coupled Plasma-Optical Emission Spectrometry (ICP-OES) experiment.

### Sample preparation and cryo-EM data acquisition

The ScBfr sample at concentration of 1.8 mg/ml was blotted (3.5□μl per blot) onto Au 300 mesh Quantifoil 1.2/1.3 grids (Quantifoil Micro Tools), and immediately plunge frozen using a Vitrobot (FEI). The grids were imaged on a Titan Krios microscope (FEI), operating at 300□kV, and equipped with a Falcon II direct detector camera (Gatan). 35-frame movies were collected at 75,000× magnification (set on microscope) in super-resolution mode with a physical pixel size of 1.11□Å/pixel (See Supplementary Material, fig S4D). The dose rate was 5.6 electrons/pixel/second, with a total exposure time of 12□s. The defocus values ranged from −0.75□μm to −2.5□μm (input range setting for data collection) as per the automated imaging software.

### Image processing

Movies were motion-corrected to compensate for the beam-induced motion using MotionCor2 (Zheng, Palovcak et al., 2017) with a B-factor of 150 pixels. All subsequent data processing was performed using cisTEM (Grant, Rohou et al., 2018) (See Supplementary Material, fig S5A). The defocus values of the motion-corrected micrographs were estimated using CTFFIND4 (Rohou & Grigorieff, 2015). The particle picking was done using auto-picking method using cisTEM and approximately, 500,000 particles were picked from the 3200 micrographs. The auto-picked particles that were subjected to multiple rounds of 2D classification to remove suboptimal particles and finally 134391particles were selected and used for further 3D classification to obtain the three models (Fig 1A-C). The 2D classification revealed various stages of the Bfr nanocages with apo and holo forms classified into various 2D classes (Fig 4B). An initial three-dimensional (3D) model was generated from our present ScBfr crystal structure and low-pass filtered to 8 Å and used as the initial reference for auto-refine. 3D reconstruction was carried out after imposing O symmetry without masking out the inner cavity/Fe-biomineral core. For the apo-ScBfr the best 77066 particles, was subjected to a final round of 3D auto-refinement and sharpen-3D to yield the final model. Similarly, for the holo-ScBfr form-I the best 19485 particles were subjected to 3D auto-refinement and sharpen-3D to yield the model of holo-ScBfr form-I. For holo-ScBfr form-II the best 13793 particles were subjected to 3D auto-refinement and sharpen-3D to yield the final model. The maps have an overall resolution of 3.4□Å for apo-ScBfr (calculated based on the gold-standard Fourier shell coefficient (FSC)□=□0.143 criterion) and 3.6Å for holo-ScBfr form-I and 4.6Å for holo-ScBfr form-II (See Supplementary Material, fig S4A-C). The angular distribution of particle projections calculated in cisTEM is shown in (See Supplementary Material, fig S5B-D). We also attempted to solve the structure without enforcing the symmetry and the subsequent 3D reconstruction model has very poor resolution and the secondary structure of the bacterioferritin were not visible in that model. The volume of the Fe-biomineral is 5% of the total volume of the nanocage cavity in the holo-ScBfr form-I but 70% in form-II. The Fe-biomineral cores inside the nanocages of the holo-ScBfr form I and II do not fit in any symmetry.

### ScBfr model building

The ScBfr crystal structure (see next section) was used as an initial model and aligned to the ScBfr cryo-EM map calculated with CisTEM. Phenix dock-in-map (Adams, Afonine et al., 2010) was used for initial model building followed by manual model building in COOT (Emsley & Cowtan, 2004) using the cryo-EM map. After initial model building, the model was refined against the EM-derived maps using the phenix.real_space_refinement tool from the PHENIX software package (Adams et al., 2010), employing rigid body, local grid, NCS, and gradient minimization. This model was then subjected to additional rounds of manual model-fitting and refinement which resulted in a final model-to-map cross-correlation coefficient of 0.83, 0.87 and 0.84 for apo-ScBfr, holo-ScBfr (form-I and form-II) respectively. Stereo-chemical properties of the model were evaluated by Molprobity (Chen, Arendall et al., 2010).

### Crystallization and structure determination

Crystallization screening was carried out at room temperature (24°C) using 1 μl of protein (10 mg/ml) mixed with equal volume of reservoir solution. Screens were setup manually by hanging-drop vapor diffusion method. All commercially available screens viz, Hampton kits, Qiagen kits and emerald biosystems kits were used for initial screening. The initial conditions were obtained from Wizard screens (Rigaku Reagents, USA). The better-quality crystals were obtained with the condition PEG3000, 0.1M HEPES pH 7.5 and 0.2 M NaCl through the grid screen optimization process. Crystals were cryo-protected in the reservoir solution supplemented with 25% glycerol and flash cooled at 100 K. The data was collected NSRRC beamline BL13B1 using ADSC Quantum 315r CCD detector at 1 Å wavelength. Data processing was done with HKL2000 program (Otwinowski & Minor, 1997). The Matthews coefficient was 2.71 Å^3^/Da (Matthews, 1968), corresponding to a solvent content of 55% with six molecules in the asymmetric unit.

The structure was determined by molecular replacement method using Phenix-Phaser program (McCoy, 2007) using the structure of bacterioferritin from *Mycobacterium smegmatis* (PDB code: 3BKN) as a search model. Initial model building was carried out using the AutoBuild program (Terwilliger, Grosse-Kunstleve et al., 2008) followed by several rounds of manual model building using COOT program (Emsley & Cowtan, 2004). The structure refinement was carried out using Phenix-Refine (Afonine, Mustyakimov et al., 2010). The model had good stereochemistry, with 99.2% residues falling within the allowed regions of the Ramachandran plot. The current model did not have well defined electron density map to build the heme molecule, instead we have observed density for modelling two Fe ions in the dimeric interface.

### DNA Library Preparation and Sequencing

A total amount of 500 ng per sample was used as input material for the DNA sample preparations. Sequencing libraries were generated using NEBNext Ultr II DNA Library Prep Kit (New England Biolabs, England) following manufacturer’s recommendations and index codes were added to attribute sequences to each sample. The genomic DNA is randomly fragmented to a size of 350 bp by Covaris cracker, then DNA fragments were end polished, A-tailed, and ligated with the full-length adapter for Illumina sequencing with further PCR amplification. At last, PCR products were purified (AMPure XP system) and libraries were analyzed for size distribution by Agilent2100 Bioanalyzer and quantified using real-time PCR.

### Inductively Coupled Plasma-Optical Emission Spectrometry (ICP-OES)

ICP-OES was used to estimate the amount of Fe in the samples. Perkin Elmer Avio 500 Inductively Coupled Plasma-Optical Emission Spectrometer was used for this study. The purified wildtype and Bfr mutants (ScBfr_1-152_ and ScBfr K42A) at a concentration of 0.01 mg/ml was subsequently treated with 0.1 M HNO_3_ to prepare the samples. Samples were analyzed on the ICP-OES along with a blank with only buffer and 0.1 M HNO_3_. Fe was not detected in the blank samples.

## Author contributions

JS and JA conceived the project; CJ, and JS designed the experiments; CJ, TYC, RR, MS, KP, performed research; TBN and SJ assisted in Cryo-EM data collection, CJ, SB and JS analysed the data; CJ and JS wrote the paper with input from other authors.

## Acknowledgments

This work was partially supported by Ministry of Education, Singapore (MoE Tier-2) grant (R-154-000-B03-112) and R154-000-A72-114 (AcRF Tier 1 grant) respectively. We acknowledge the NSRRC, Taiwan beamline 13B1 for crystallographic data collection and NUS CBIS for cryo-EM data collection. We thank Dr Sujatha Narayanankutty, Department of Biological Sciences, National University of Singapore (NUS) for advice in DNA sequencing studies, Dr Kang Wei Tan, Department of Chemistry, NUS and Dr Matthew Belousoff, Monash University, Australia, for the discussions on EM experiments and analysis.

## Data availability

The coordinates of the ScBfr crystal structure was deposited in wwPDB under the accession number 5XX9. The coordinates of the ScBfr nanocage structures and the cryo-EM density map are deposited in wwPDB and EMDB under accession number 6K3O, EMD-9910 (apo-form), 6K43, EMD-9913 (holoform-I), 6K4M, EMD-9915 (holoform-II).

